# Genome-wide association analyses in dairy heifers highlight genes overlapping with mouse and human fertility and human health traits

**DOI:** 10.1101/2024.12.19.629410

**Authors:** Mackenzie A. Marrella, Gustavo P. Schettini, Michael Morozyuk, Allison Walsh, Rebecca Cockrum, Fernando H. Biase

## Abstract

Heifer Infertility and disease are important challenges in dairy cattle production. We investigated genetic differences between Holstein heifers with varying fertility potential and health. We carried out a genome-wide association analysis comparing heifers that conceived at first insemination against those requiring multiple attempts or failing to become pregnant, as well as heifers culled due to health issues. There were 12 significant SNPs (P<5x10^-5^) associated with fertility and 35 SNPs associated with health traits. There were 166 significant SNPs when infertile, sub-fertile and animals culled due to health issues were grouped. Two SNPs identified in the analysis of infertility were found near *NUFIP1* and within *TENM4* genes, both genes are linked to embryonic lethality in mouse knockouts. Follow-up CRISPR-Cas9 mediated disruption of *NUFIP1* significantly (P<0.05) reduced *in vitro* blastocyst development in cattle embryos, while *TENM4* editing did not alter *in vitro* blastocyst development. Additionally, SNPs overlapped with previously identified reproduction-related QTL (*CNTN4*, *DLG2*, *PARP10*, *PRICKLE*, *TMEM150B*) or health-related QTL (*FAM162A*, *PARP10*). We also identified genes within or near genes previously associated with age at menarche (*CADM2, DLG2*, *FHIT*, *LSAMP* and *TENM4*) or lung function or pulmonary diseases (*ASCC2*, *BCAS3*, *BTBD9*, *CADM2*, *CNTN4*, *CPEB4*, *CTNNA2*, *DEUP1*, *DGKH*, *DLG2*, *ENOX1*, *EPHB1*, *ERC2*, *ERGIC1*, *EYA2*, *FAM162A*, *FGF18*, *FHIT*, *GRID1, KCNIP4*, *LINGO2*, *LRMDA*, *MALRD1*, *NEBL*, *PLA2G6*, *PLXDC2*, *PRPF18*, *SLC8A1*, *TEAD4*, *TSPAN9*) in humans. These results further support genetic components of fertility and health in cattle. The findings also show overlapping genetic architecture between fertility and health traits, with a degree of conservation across mammals.

**Summary sentence:** Several genetic variants that influence female fertility and health in cattle were identified, and many genes harboring or near significant polymorphisms are common to equivalent phenotypes in mice and humans.

## Introduction

The sustainability of dairy farms depends on having as many healthy cows in lactation as possible. Therefore, the constant replacement of cows that leave the herd due to culling or death is essential for sustainable dairy cattle production. Most small (67.7%) and medium (69.2%) dairy operations raise all heifers on-site, but 40% of the large operations raise heifers on-site (USDA–APHIS–VS–CEAH, September 2021). Raising replacement heifers is the second largest source of expenses for dairy farms, only second to feeding costs[1]. Therefore, producers need to implement careful management strategies to avoid severe financial losses that cannot be recovered. Losses associated with heifer development are mostly incurred due to infertility (∼5% [2]) and death (7.8% and 1.8% pre-weaning and post-weaning, respectively [3]).

Infertility, or subfertility, and disease are major barriers to the profitability and sustainability of dairy cattle production in the United States [4]. Heifer fertility is particularly important to producers because heifers that calve at the appropriate time have increased productivity and longevity in the herd[1, 5–7]. In parallel, disease in heifers has also been associated with decreased productivity. Heifers that are affected by disease have decreased milk production in their first lactation[8], increased calving intervals[9], and increased age at first calving [10]. On the more extreme end of the spectrum, heifers may die from diverse diseases, with a greater prevalence of respiratory-related infections [3]. Collectively, infertility and death by disease are the major reasons preventing heifers from contributing to a productive dairy herd. Therefore, being able to identify these heifers earlier in their life will help to improve the sustainability of dairy operations.

Molecular genotyping is a promising approach for early decision-making in heifer management [11]. Several SNPs have been significantly associated with dairy heifer-related traits, such as conception rate [2, 12–16], early calving [2, 17], times inseminated to achieve pregnancy [2], inability to achieve pregnancy after multiple artificial inseminations [18], or multiple-traits analysis [19]. Likewise, SNPs have been significantly identified for heifer livability [17]. While progress has been made in the identification of genomic variants significantly associated with traits that are relevant to heifer development, limited effort has been devoted to the identification of causal mutations or genes. The two examples of such studies were carried out by Ortega et al. [20]and Adams et al. [21] that identified mutations in the genes Intraflagellar Transport 80 (*IFT80*) and Apoptotic Peptidase Activating Factor 1 (*APAF1*), respectively, leading to embryonic losses in dairy cattle.

Here, we carried out a study to test the hypothesis that there are systemic genetic differences between (a) Holstein heifers of different fertility potential and (b) heifers that remain in the herd as replacements or leave the herd before 13 months of age due to health reasons. The objectives of this study were to identify genomic variants associated with heifer fertility or health in Holsteins. Two of the genomic variants significantly associated with heifer infertility guided a follow-up hypothesis-driven mechanistic experiment to test if the disruption of the genes *NUFIP1* and *TENM4* have a lethal effect on cattle pre-implantation embryo development.

## Material and Methods

Ethics: No live animal was handled for this research.

### Data collection

We obtained genomic data from the Virginia Tech Dairy Center from The Council on Dairy Cattle Breeding (CDCB). This dataset contained 78,964 genotypes from 746 Holstein heifers. We also obtained reproductive and health data from the Virginia Tech Dairy Center using the herd management software PCDART encompassing the years 2007-2023. Data were filtered to retain only Holstein heifers that had complete records of phenotype and health-related information.

### Processing of genotypes for analysis

We recorded genotypes as missing (00), homozygous reference allele (11), heterozygous (12), and homozygous alternate allele (22) in R software (v.4.2.3) [22]. Then, the data was filtered to include only records for Holstein heifers that had phenotypes and genotypes. After filtering, 283 animals were retained for further analysis. From this data, MAP and PED files were created for analysis in PLINK (v1.90b6.18) [23, 24]. SNPs were then removed if they had a minor allelic frequency < 0.01, a genotypic frequency < 0.05, or deviated from Hardy-Weinberg Equilibrium (*P* < 0.00001). Finally, to identify population stratification in our sample, we conducted a principal component analysis in PLINK. Results were visualized using the package “ggplot2” [25] in R software.

### Genome-wide association analyses

We separated the heifers into four groups (Fig. 1A): (a) delivered a healthy calf after their first artificial insemination (N=271); (b) delivered a healthy calf after four or five artificial inseminations (N= 28); (c) did not deliver a calf after four or more artificial inseminations and were removed from the herd due to reproductive reasons (N=11); and (d) were removed from the herd before 13 months of age due to health reasons (N=14).

We conducted association analyses with heifers that delivered a healthy calf after their first artificial insemination as a control group, comparing it with other groups to test specific hypotheses. We used Fisher’s exact test in PLINK to evaluate allelic, dominance, and recessive associations. Associations were considered significant at alpha = 5x10^-5^ [2, 26–28].

### Databases and approaches for obtaining biological insights

We used the “GALLO” [29]R package to annotate significant SNPs based on the following databases: Ensembl [30–32], and Animal QTLdb[33–35]. Genes were screened for overlap with pertinent mouse knockout phenotypes [36–38] as well as pertinent phenotypes previously investigated in human GWAS studies, compiled in the National Human Genome Research Institute-European Bioinformatics Institute GWAS catalog[39]. In addition, we used the “topGO” R package to conduct a Fisher’s exact test [40, 41] to evaluate Gene Ontology [42, 43] enrichment [40] on a set of genes associated with significant SNPs while using protein-coding genes and long non-coding genes in the Ensembl annotation as a background.

### DNA editing of targeted genes in cattle embryos

We designed guide RNAs using the CRISPOR [75]. All gRNAs were purchased as a single guide RNA molecule (sgRNA) comprising crRNA and transacting crRNA (tracerRNA) from IDT (Integrated DNA Technologies, Research Triangle Park, NC, USA), as well as CRISPR-Cas9D10A nickase V3. The gRNA sequences to edit the gene *NUFIP1* were 5’-GATAACGGAAGTGACGCCTAAGG-3’ and 5’ATCACAAGCTGATATGTCATCGG-3’, and for the gene *TENM4* were 5’-GAGGAACTTGCACCCAAAGT-3’ and 5’-CTGTGCCACTTACGGCTGCG-3’.

*In vitro* embryo production and DNA editing were conducted according to protocols, including media composition, detailed elsewhere. [44–46]. We started with the collection of cumulus-oocyte complexes (COCs) obtained from ovaries sourced from a slaughterhouse. The COCs were selected based on morphology and cultured in oocyte maturation medium in groups of 10 in 50 µL of medium covered by light mineral oil. In vitro maturation plates were incubated for 22-24 hours at 38.5 °C and 5 % CO_2_ in a humidified atmosphere. Next, we moved the mature COCs through synthetic oviductal fluid (SOF) medium, containing N-2-hydroxyethylpiperazine-Nʹ-2-ethanesulfonic acid (HEPES; Thermo Fisher Scientific) and SOF for fertilization before transferring into a final fertilization plate (100 COCs/ml). We prepared sperm from thawed frozen semen straws and processed for in vitro fertilization at a concentration of 1,000,000 spermatozoa/ml. COCs and spermatozoa were co-incubated for 12–13 hours under the same conditions described for in vitro maturation.

We mixed CRISPR-Cas9D10A and sgRNAs for the formation of ribonucleoproteins in OptiMEM reduced serum medium (Thermo Fisher Scientific, Grand Island, NY) at room temperature for at least one hour before electroporation. The concentrations in the solution for the formation of RNPs were 800ng/µl Cas9D10A and 800ng/µl of each sgRNA.

We removed the cumulus cells from the PZs by pulsed vortexing in a hyaluronidase solution followed by repeated pipetting and electroporated them in Opti-MEM media containing RNPs at the concentration of 400ng/µl Cas9D10A and 400ng/µl of each sgRNA. The electroporation parameters were as follows: six poring pulses of 15V, with 10% decay, for 2ms with a 50ms interval, immediately followed by six transfer pulses of 3V, 40% decay, for 50ms with a 50ms interval, alternating the polarity. We conducted two electroporation sessions, the first at 14 hours post fertilization (hpf) and the second at 20 hpf. After the second electroporation, PZs were placed in culture media (25 in 50 µl of synthetic oviduct fluid), covered by mineral oil, and incubated as indicated above 38.5 °C with 5 % CO_2_, 5 % O_2_, and 90 % N_2_ in a humidified Eve Benchtop Incubator (WTA, College Station, TX, USA).

For each target DNA, we executed the experiment in triplicate. We recorded the number of embryos that cleaved at ∼45 hpf and blastocysts at ∼168 hpf and ∼190 hpf. For statistical analysis, we considered culture drops to be biological replicates. We analyzed count data (success of blastocyst development or developmental arrest) using a general linear model with a binomial family, which results in logistic regression analysis [47] using the “glmer” function from the R package “lme4” [48]. We used the number of blastocysts and the number of putative zygotes that failed to develop into blastocysts as the dependent variable. Group (Cas9 + targeting gRNAs or Cas9 + scramble gRNAs) was a fixed effect, and replicate was a random variable. The Wald statistical test [49] was conducted with the function “Anova” from the R package “car”. Finally, we carried out a pairwise comparison using the odds ratio and two-proportion z-test and t-test [50] employing the “emmeans” function of the R package “emmeans”. The null hypothesis assumed that the odds ratio of the proportion (*p*) of the two groups was not different from 1 (*H*_0_: *p*_1_/*p*_2_ = 1). We inferred significance when adjusted P value < 0.05.

To evaluate the edits, we collected embryos at the 8-cell stage or morula five days post-culture. First, we removed their zona pellucida with EmbryoMax Acidic Tyrode’s Solution (Millipore Sigma, Danvers, MA), washed the embryos extensively in phosphate buffer saline, and placed embryos individually in a microcentrifuge tube. We exposed their DNA by adding four µL of QuickExtract DNA Extraction Solution (Biosearch Laboratories, USA) and incubating the solution at 65°C for 15 min followed by 2 min at 98°C and holding at 4°C. Next, we conducted PCR reactions using the following oligonucleotides designed using NCBI’s Primer-BLAST [51] (*NUFIP1*: 5’ ACTGCAAATCCCAGGGCTAC 3’, 5’ AACATCACAGGATGGCAGGG 3’; *TENM4*: 5’ GCTCTGGGTTATTGTGGCCT 3’, 5’ AGGCAGGGGATAAAACAGCC 3’). In addition to the cellular extract, the PCR reaction mix for the *NUFIP1* gene segment consisted of 0.2 IU/μL Phusion Hot Start II DNA Polymerase (Thermo Fisher Scientific), 1× Phusion HF Buffer, 200 μM dNTPs (Promega, Madison, WI, USA), and forward and reverse oligonucleotides (IDT, Coralville, IA, USA) at 0.1 μM each, in a final volume of 50 μL in 0.2 ml PCR tubes. The PCR reaction mix for the *TENM4* fragment consisted of PrimeSTAR® GXL DNA Polymerase (Takara Bio Inc) 1× PrimeSTAR Buffer, 200 μM dNTPs, and forward and reverse oligonucleotides at 0.1 μM each, in a final volume of 50 μL in 0.2 ml PCR tubes. The thermocycling conditions were: 98°C for 1 min, followed by 35 cycles of 98°C for 15 s, 60°C for 30 s, and 72°C for 3 min, followed by a final extension of 4 min at 72°C.

We confirmed the amplification by assaying 5 µL of each amplicon by electrophoresis on a 1.5% Agarose gel before staining with Diamond Nucleic Acid Dye (Promega, Madison, WI) and imaging. Finally, we prepared libraries for amplicon sequencing with the Native Barcoding Kit 96 V14 (Oxford Nanopore Technologies, Lexington, MA, USA), following the manufacturer’s instructions. We sequenced the libraries in a MinION flowcell (R10.4.1) using a MinION Mk1C (Oxford Nanopore Technologies, Lexington, MA, USA) following the manufacturer’s recommendations. We carried out super accuracy base calling with Guppy v6.4.2 [52] followed by quality filtering using Fitlong (https://github.com/rrwick/Filtlong) to remove sequences < 2000 nucleotides long and mean quality less than 80% accuracy. Next, we mapped filtered sequences to the cattle reference genome from Ensembl (ARS-UCD1.2) using minimap2 v2.27[53, 54], allowing for spliced alignment, and only primary alignments were retained with SAMtools [55, 56]. Next, we removed sequences that contained insertions greater than 50 nucleotides and those that required soft clipping of more than 150 nucleotides for alignment. Finally, we only retained alignments with less than 2% of nucleotide mismatches.

## Results

### Overview of the data and groups evaluated

We matched genotype and phenotype data for 324 Holstein heifers, which we separated into four groups based on their health and fertility (Fig. 1A). An overview of the genotypic data (78,964 SNPs) showed no clustering of samples due to genetic structure or independent clustering that should be considered for the hypothesis testing (Fig. 1B).

**Fig. 1.**
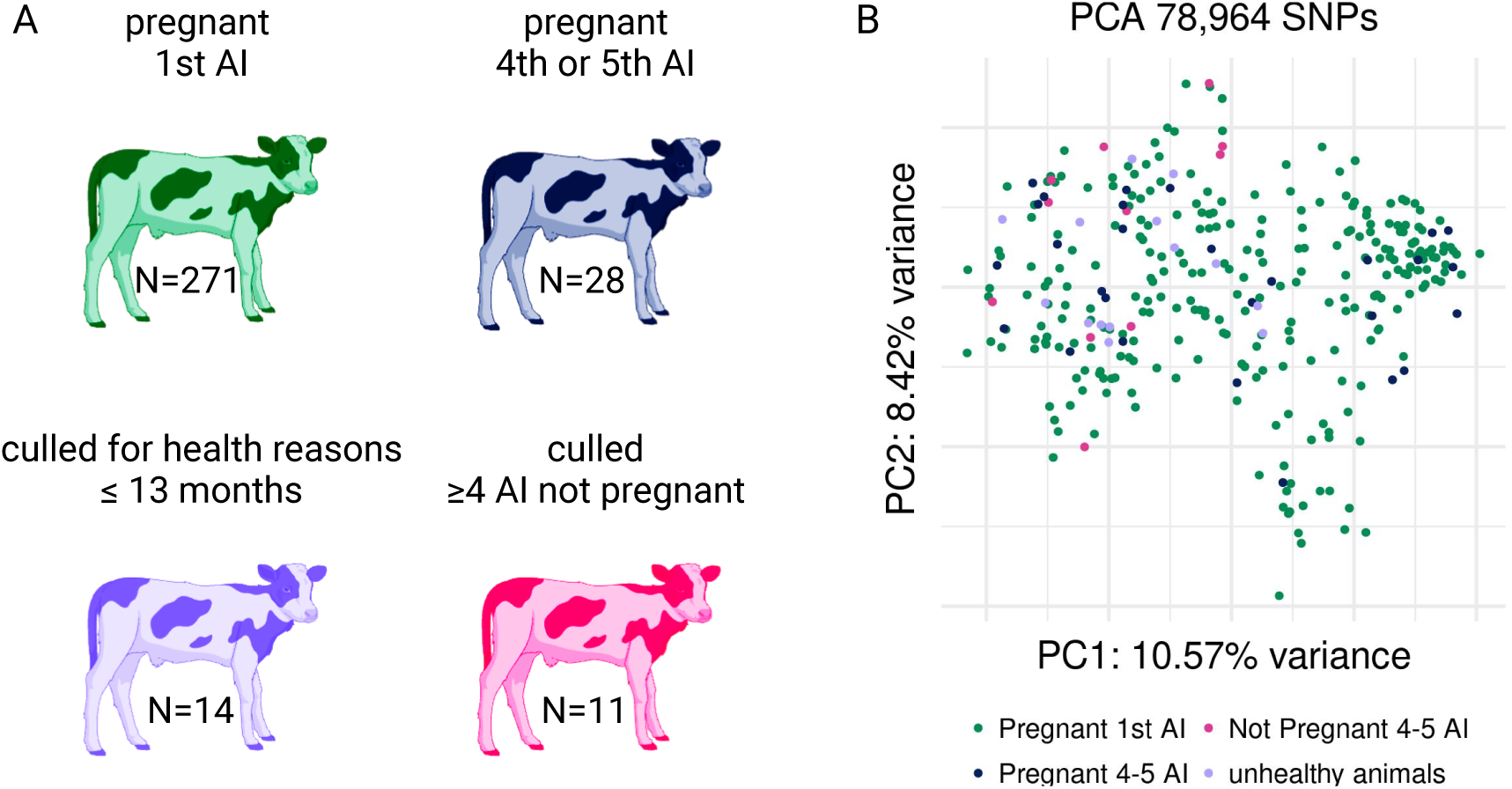
Overview of the data. (A) Sample sizes for each group used for hypothesis testing. (B) Principal component analysis of the genotypes for 324 samples utilized in the studies.

### Genome-Wide Association Analysis

In order to identify SNPs that were associated with infertility in heifers, we compared the heifers that were pregnant to the first artificial insemination to the heifers who were artificially inseminated for two or more services and failed to get pregnant. There were two SNPs (rs110887612 and rs41618427) with allelic frequency and dominance association with infertility, and three SNPs (rs137239652, rs135984630 and rs110392437) with dominance relationship of the alleles associated with infertility. We also tested allelic and genotypic associations with heifers that were pregnant after four or five artificial insemination attempts. The SNPs rs3423415998, rs135730562, and rs109809439 showed allelic frequency and dominance relationship of alleles associated with the number of artificial inseminations, while SNPs rs109014975, rs41618427, rs110123517, rs110354925, rs133314893 showed a dominance relationship of alleles associated with number of artificial inseminations needed to achieve a pregnancy and delivery a calf (Table 1). Four SNPs (rs110887612, rs110392437, rs109014975, rs110354925) were located within gene boundaries (*TRIM65*, *TENM4*, *FECH*, *TBC1D1*, respectively), and the SNP rs41618427 is 9,560 nucleotides downstream of the gene *NUFIP1* (Table 1).

**Table 1.** Summary of association analysis for heifer fertility.

| Variant identifier | Position | P-value |  | Gene | Nearest gene |
| --- | --- | --- | --- | --- | --- |
|  |  | Fisher's exact test | Dom / Rec |  |  |
| pregnant to the 1<sup>st</sup> AI vs. those that did not get pregnant |  |  |  |  |  |
| rs110887612 | 19:55732113 | 2.31x10<sup>-5</sup> | 2.98x10<sup>-5</sup> | TRIM65<sup>ab3UTR</sup> | - |
| rs41618427 | 12:15108440 | 2.72x10<sup>-5</sup> | 2.47x10<sup>-5</sup> | - | NUFIP1 |
| rs137239652 | 16:23978490 | ns | 2.47x10<sup>-5</sup> | - | - |
| rs135984630 | 29:15706087 | ns | 3.08x10<sup>-5</sup> | - | - |
| rs110392437 | 29:16796262 | ns | 4.27x10<sup>-5</sup> | TENM4<sup>abi</sup> | - |
| pregnant to the 1<sup>st</sup> AI vs. those that needed 4 or 5 AI before carrying a pregnancy to term |  |  |  |  |  |
| rs3423415998 | AI:20092858 | 2.88x10<sup>-6</sup> | 2.21x10<sup>-6</sup> | - | - |
| rs135730562 | 17:20107776 | 4.74x10<sup>-6</sup> | 3.82x10<sup>-6</sup> | - | - |
| rs109809439 | 20:16527323 | 3.05x10<sup>-5</sup> | 3.37x10<sup>-5</sup> | - | - |
| rs109014975 | 24:56805748 | ns | 2.35x10<sup>-5</sup> | FECH<sup>abi</sup> | - |
| rs41618427 | 12:15108440 | ns | 2.63x10<sup>-5</sup> | - | NUFIP1 |
| rs110123517 | 17:3325010 | ns | 3.85x10<sup>-5</sup> | - | - |
| rs110354925 | 6:57353195 | ns | 4.72x10<sup>-5</sup> | TBC1D1<sup>abi</sup> | - |
| rs133314893 | 13:60180672 | ns | 4.78x10<sup>-5</sup> | - | - |

ns = not significant; <sup>a</sup> annotated in the NCBI database; <sup>b</sup> annotated in the Ensemble database; <sup>i</sup> intron; <sup>3UTR</sup>
three prime untranslated region; - intergenic region.

We also compared the genotypes between heifers that were pregnant after the first artificial insemination to those who left the herd before 13 months of age due to health reasons (Table 2). The allelic association test identified one SNP with a strong association with health rs42250917 and 18 SNPs with moderate association for the test of allelic frequencies. There were 28 SNPs with a dominance association with the heifer health (Table 2). Sixteen out of the 35 SNPs were located within gene boundaries (Table 2).

**Table 2.**
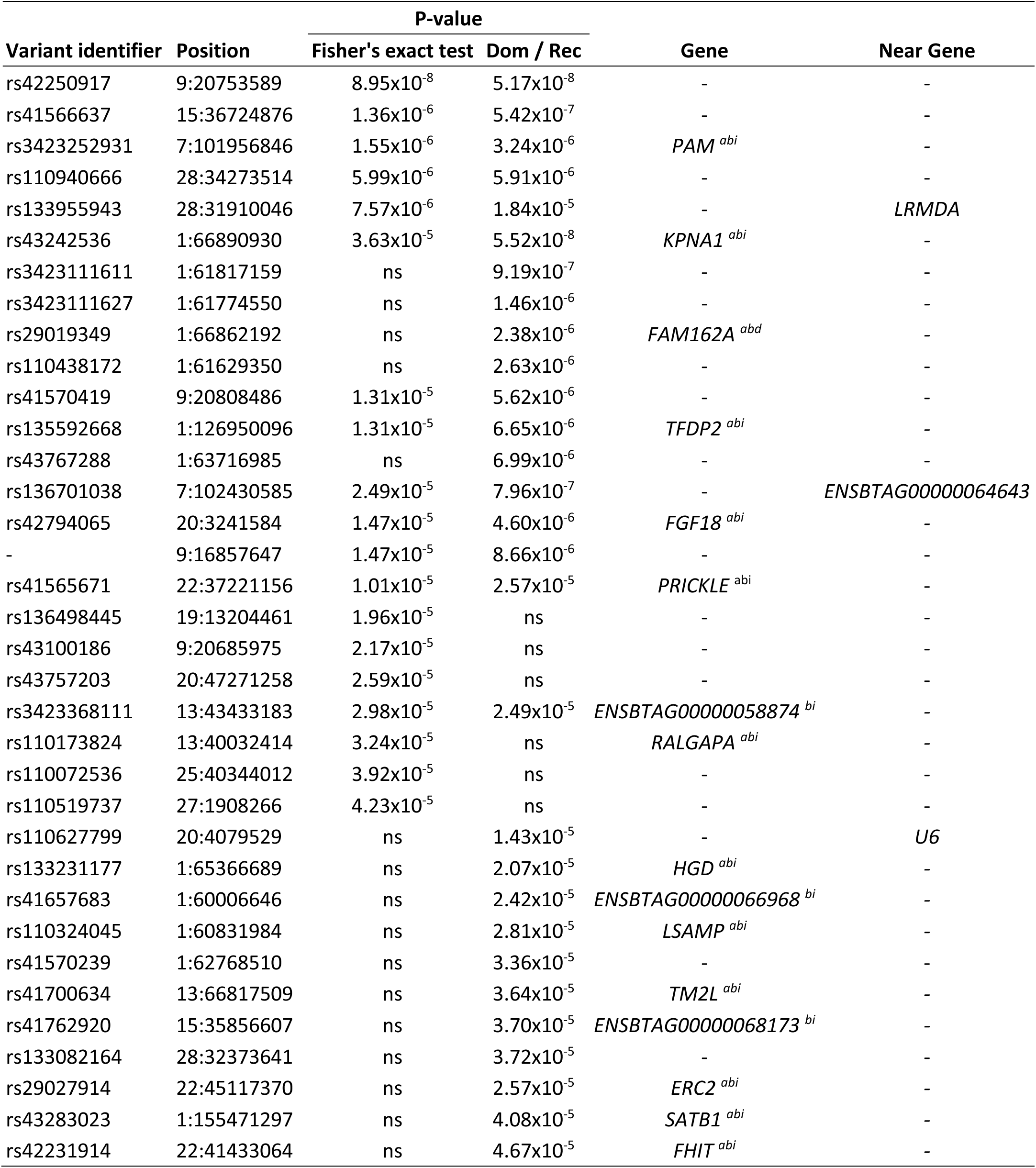
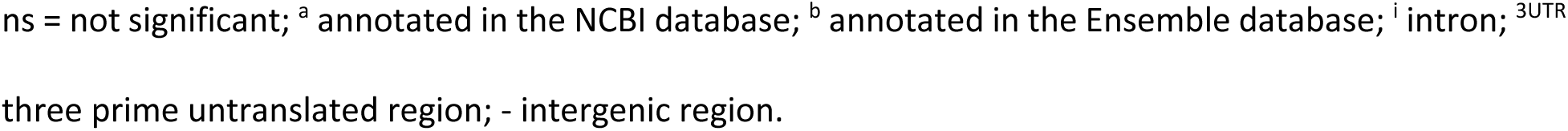
Summary of association analysis for heifer health.

| Variant identifier | Position | P-value |  | Gene | Near Gene |
| --- | --- | --- | --- | --- | --- |
|  |  | Fisher's exact test | Dom / Rec |  |  |
| rs42250917 | 9:20753589 | 8.95x10<sup>-8</sup> | 5.17x10<sup>-8</sup> | - | - |
| rs41566637 | 15:36724876 | 1.36x10<sup>-6</sup> | 5.42x10<sup>-7</sup> | - | - |
| rs3423252931 | 7:101956846 | 1.55x10<sup>-6</sup> | 3.24x10<sup>-6</sup> | *PAM*<sup>abi</sup> | - |
| rs110940666 | 28:34273514 | 5.99x10<sup>-6</sup> | 5.91x10<sup>-6</sup> | - | - |
| rs133955943 | 28:31910046 | 7.57x10<sup>-6</sup> | 1.84x10<sup>-5</sup> | - | *LRMDA* |
| rs43242536 | 1:66890930 | 3.63x10<sup>-5</sup> | 5.52x10<sup>-8</sup> | *KPNA1*<sup>abi</sup> | - |
| rs3423111611 | 1:61817159 | ns | 9.19x10<sup>-7</sup> | - | - |
| rs3423111627 | 1:61774550 | ns | 1.46x10<sup>-6</sup> | - | - |
| rs29019349 | 1:66862192 | ns | 2.38x10<sup>-6</sup> | *FAM162A*<sup>abd</sup> | - |
| rs110438172 | 1:61629350 | ns | 2.63x10<sup>-6</sup> | - | - |
| rs41570419 | 9:20808486 | 1.31x10<sup>-5</sup> | 5.62x10<sup>-6</sup> | - | - |
| rs135592668 | 1:126950096 | 1.31x10<sup>-5</sup> | 6.65x10<sup>-6</sup> | *TFDP2*<sup>abi</sup> | - |
| rs43767288 | 1:63716985 | ns | 6.99x10<sup>-6</sup> | - | - |
| rs136701038 | 7:102430585 | 2.49x10<sup>-5</sup> | 7.96x10<sup>-7</sup> | - | *ENSBTAG00000064643* |
| rs42794065 | 20:3241584 | 1.47x10<sup>-5</sup> | 4.60x10<sup>-6</sup> | *FGF18*<sup>abi</sup> | - |
| - | 9:16857647 | 1.47x10<sup>-5</sup> | 8.66x10<sup>-6</sup> | - | - |
| rs41565671 | 22:37221156 | 1.01x10<sup>-5</sup> | 2.57x10<sup>-5</sup> | *PRICKLE*<sup>abi</sup> | - |
| rs136498445 | 19:13204461 | 1.96x10<sup>-5</sup> | ns | - | - |
| rs43100186 | 9:20685975 | 2.17x10<sup>-5</sup> | ns | - | - |
| rs43757203 | 20:47271258 | 2.59x10<sup>-5</sup> | ns | - | - |
| rs3423368111 | 13:43433183 | 2.98x10<sup>-5</sup> | 2.49x10<sup>-5</sup> | *ENSBTAG00000058874*<sup>bi</sup> | - |
| rs110173824 | 13:40032414 | 3.24x10<sup>-5</sup> | ns | *RALGAP*<sup>abi</sup> | - |
| rs110072536 | 25:40344012 | 3.92x10<sup>-5</sup> | ns | - | - |
| rs110519737 | 27:1908266 | 4.23x10<sup>-5</sup> | ns | - | - |
| rs110627799 | 20:4079529 | ns | 1.43x10<sup>-5</sup> | - | *U6* |
| rs133231177 | 1:65366689 | ns | 2.07x10<sup>-5</sup> | *HGD*<sup>abi</sup> | - |
| rs41657683 | 1:60006646 | ns | 2.42x10<sup>-5</sup> | *ENSBTAG00000066968*<sup>bi</sup> | - |
| rs110324045 | 1:60831984 | ns | 2.81x10<sup>-5</sup> | *LSAMP*<sup>abi</sup> | - |
| rs41570239 | 1:62768510 | ns | 3.36x10<sup>-5</sup> | - | - |
| rs41700634 | 13:66817509 | ns | 3.64x10<sup>-5</sup> | *TM2L*<sup>abi</sup> | - |
| rs41762920 | 15:35856607 | ns | 3.70x10<sup>-5</sup> | *ENSBTAG00000068173*<sup>bi</sup> | - |
| rs133082164 | 28:32373641 | ns | 3.72x10<sup>-5</sup> | - | - |
| rs29027914 | 22:45117370 | ns | 2.57x10<sup>-5</sup> | *ERC2*<sup>abi</sup> | - |
| rs43283023 | 1:155471297 | ns | 4.08x10<sup>-5</sup> | *SATB1*<sup>abi</sup> | - |
| rs42231914 | 22:41433064 | ns | 4.67x10<sup>-5</sup> | *FHIT*<sup>abi</sup> | - |

ns = not significant; <sup>a</sup> annotated in the NCBI database; <sup>b</sup> annotated in the Ensemble database; <sup>i</sup> intron; <sup>3</sup>UTR
three prime untranslated region; - intergenic region.

Next, we carried out an analysis that included data on healthy heifers and those heifers that were culled from the herd for health or reproductive reasons. There were 166 SNPs with moderate to strong association between the two groups, 137 and 95 significant for the test of allelic frequencies and dominance, respectively. Interestingly, 94 of the significant SNPs were within the boundaries of 82 genes (Table 3).

**Table 3.**
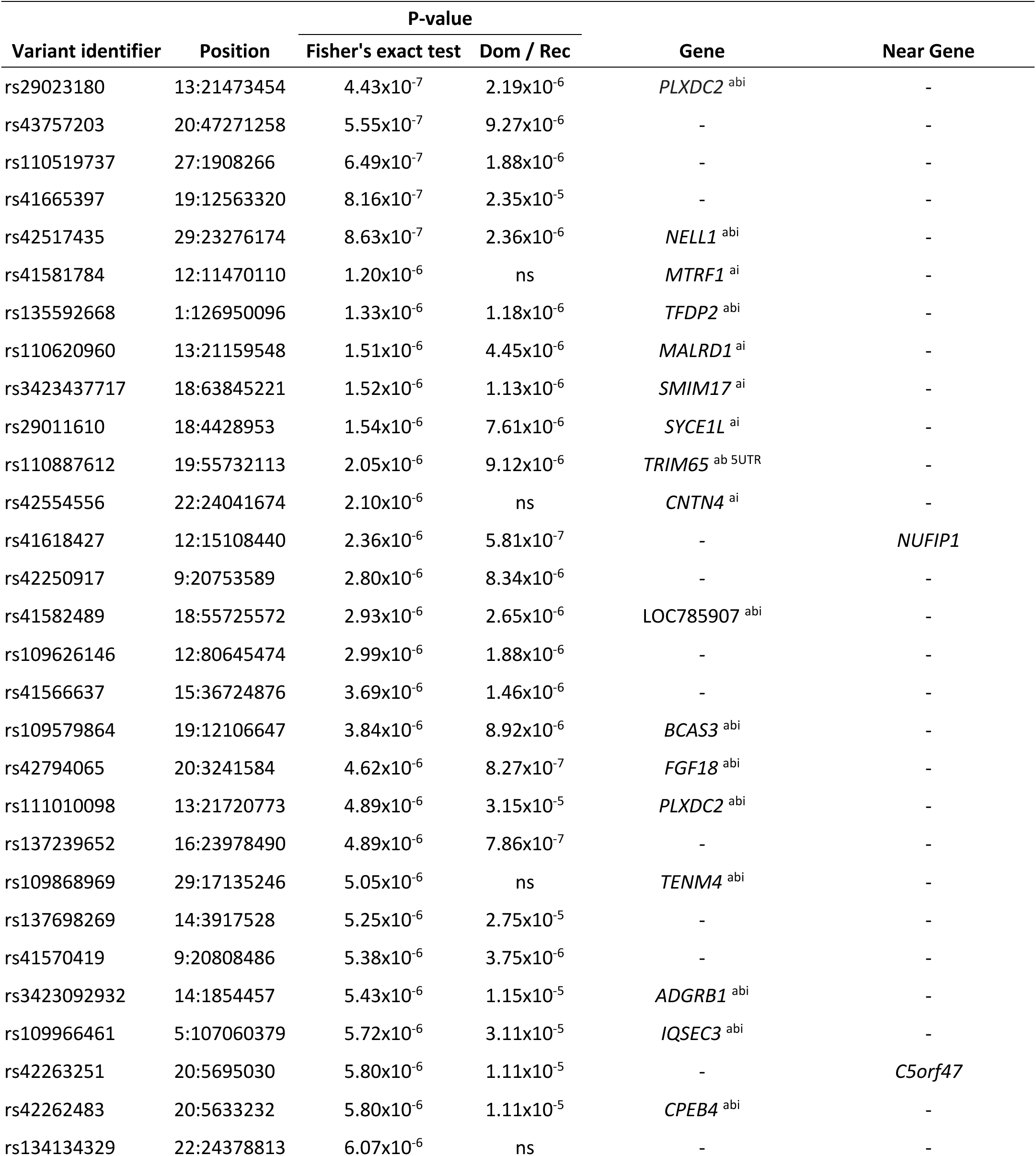
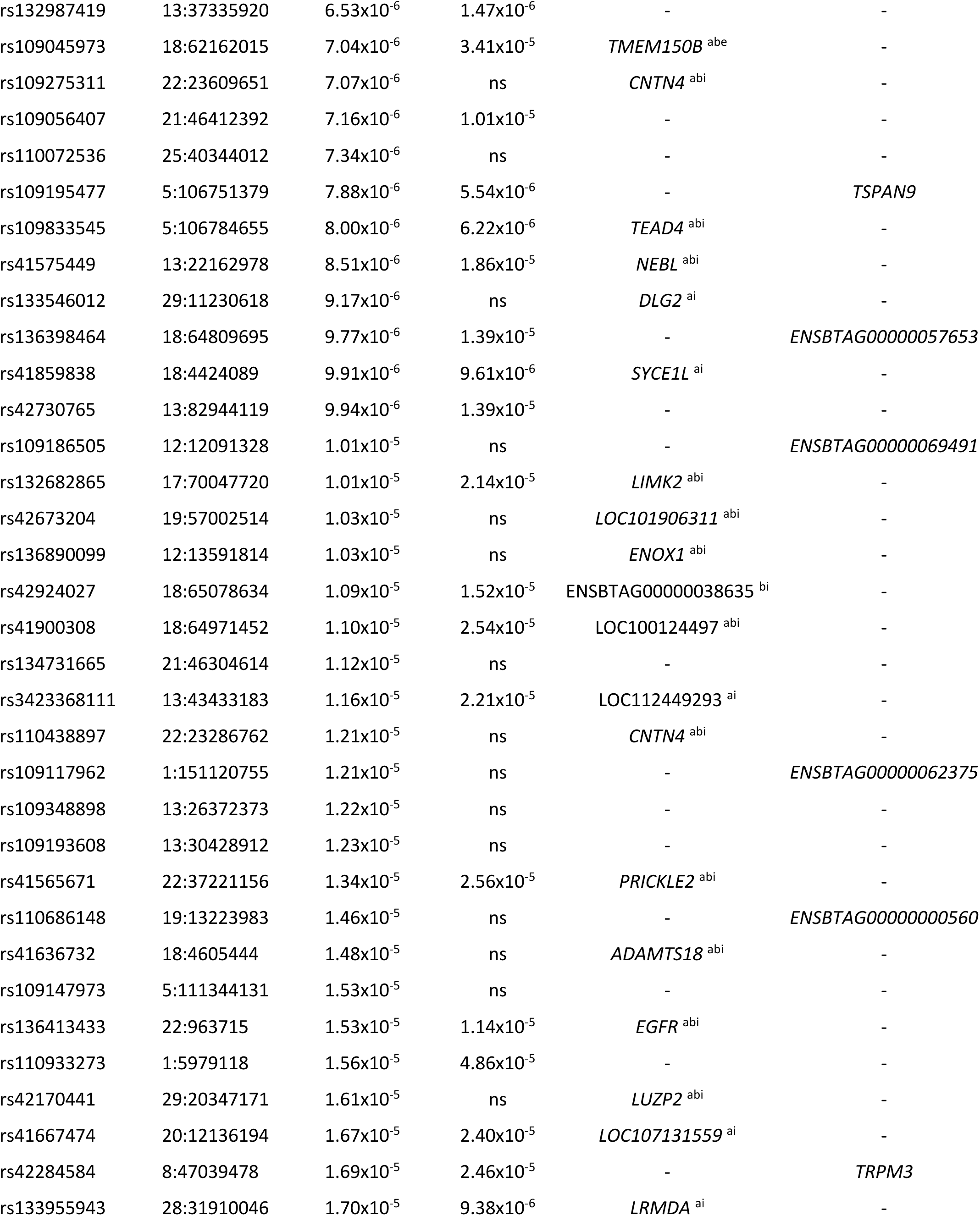
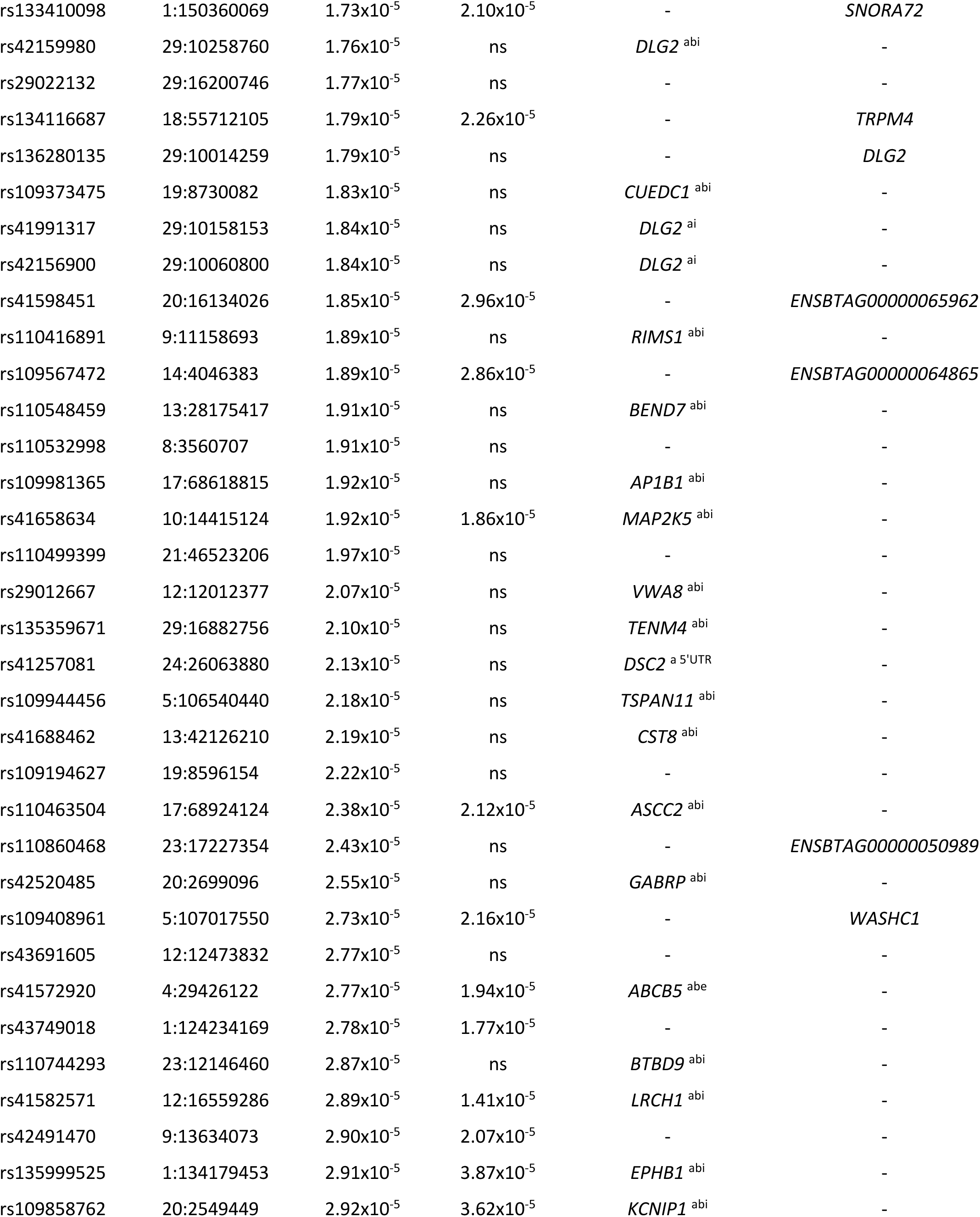
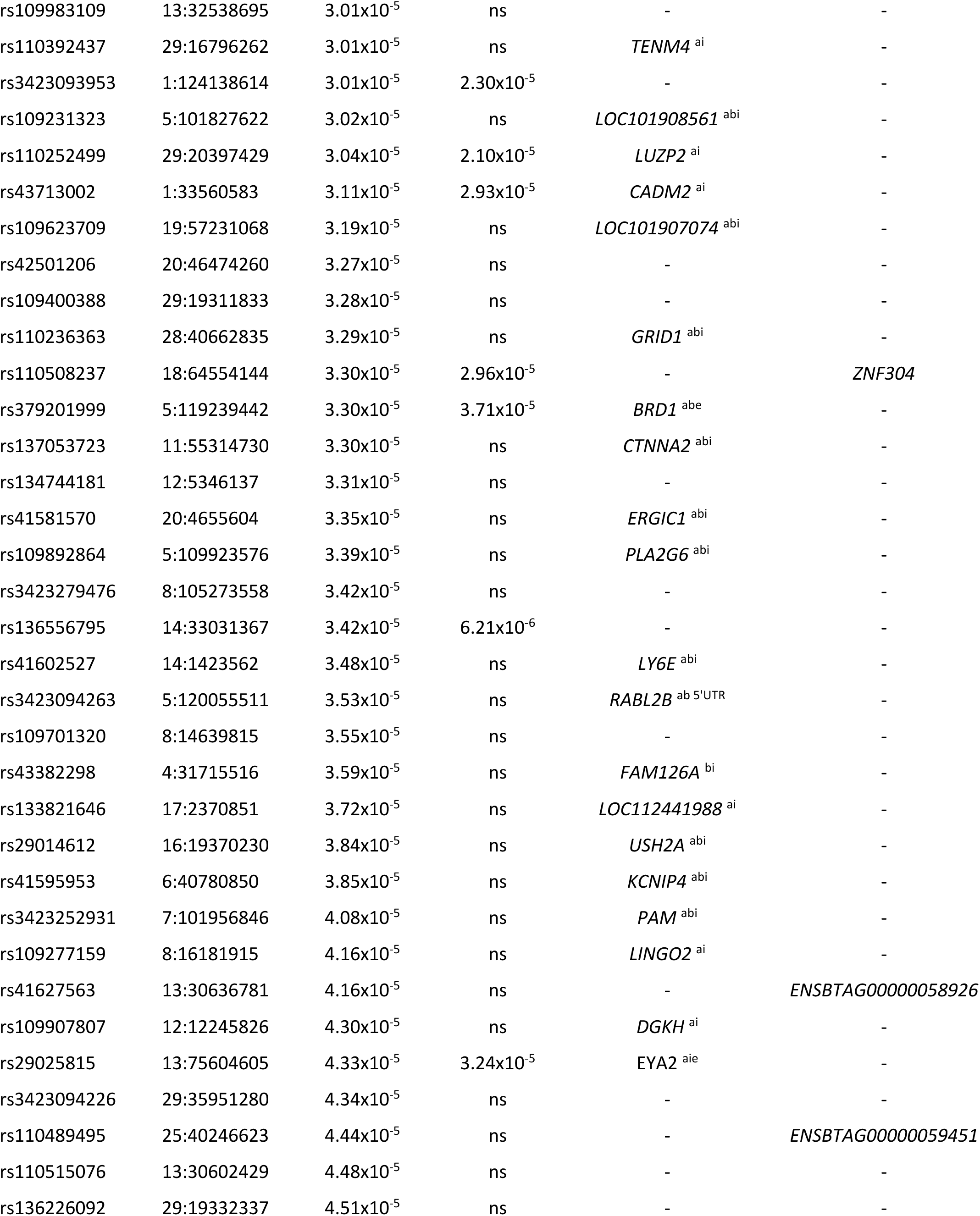
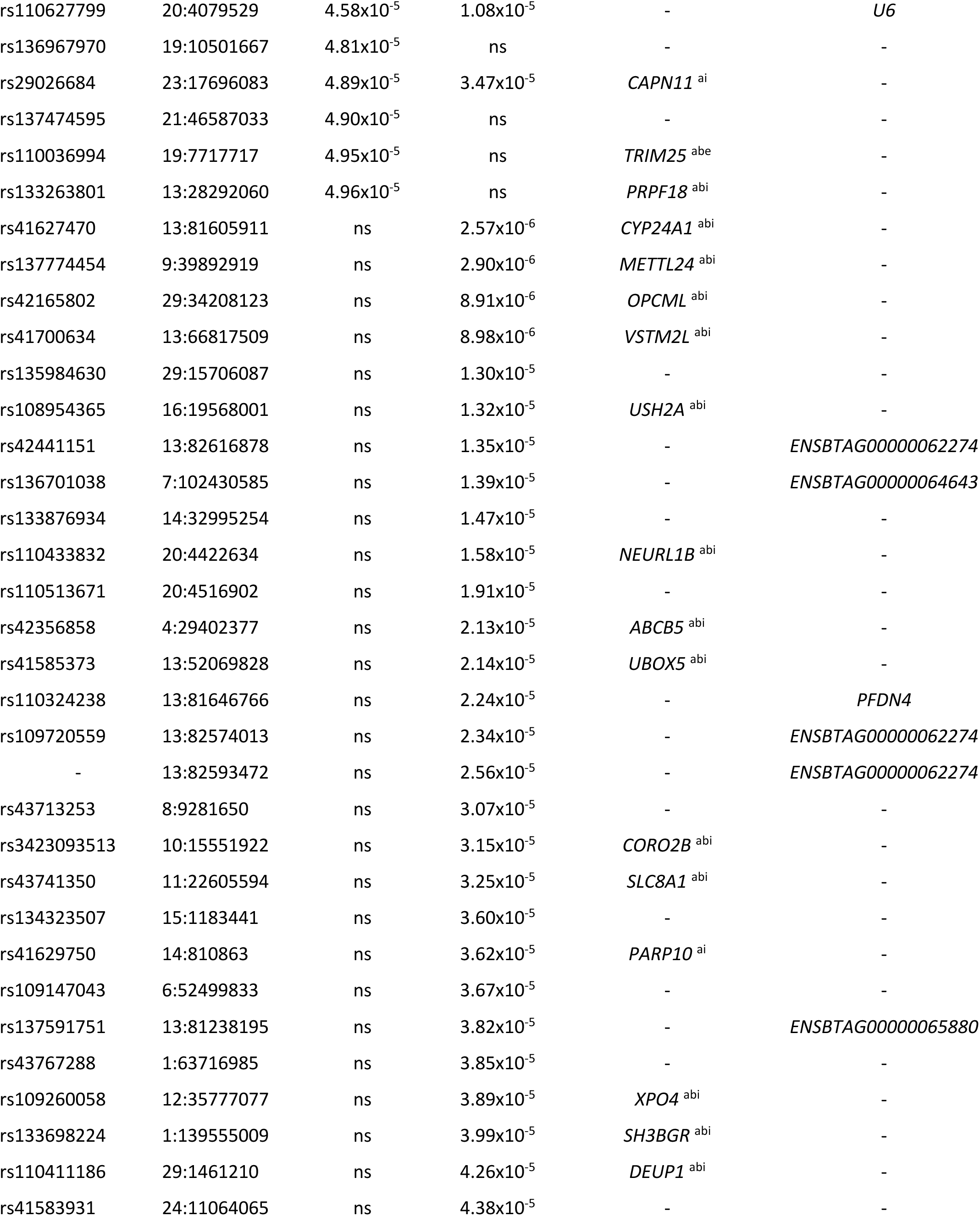
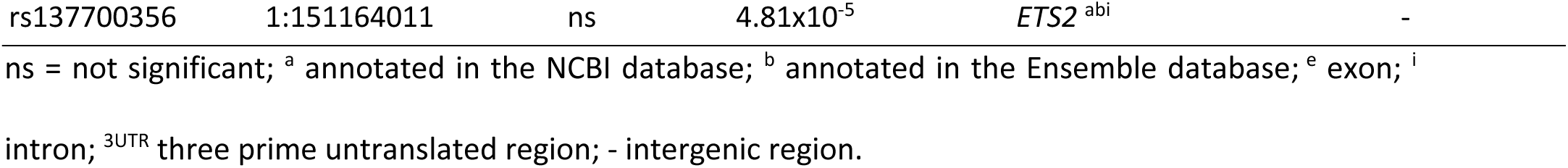
Summary of association analysis for heifers that were culled from the herd due to infertility and health.

### In silico annotation of significant SNPs and associated genes

We annotated significant SNPs and corresponding genes, if available (Fig. 2). There were seven genes (*BTBD9*, *DLG2*, *ERC2*, *GABRP*, *GRID1*, *KPNA1*, *RIMS1*) related to 11 SNPs that composed a significantly enriched biological process “synaptic signaling” (FWER<0.05). Thirty-four SNPs within or near six genes were located within health-related QTL (*FAM162A* [57], *PARP10* [58]) or reproduction-related QTL (*CNTN4*, *DLG2*, *PARP10*, *PRICKLE*, *TMEM150B*). There were 17 SNPs at or near 15 genes (*ETS2*, *ASCC2*, *BCAS3*, *BRD1*, *CST8*, *EGFR*, *LY6E*, *MAP2K5*, *NUFIP1*, *PRPF18*, *SLC8A1*, *TEAD4*, *TENM4*, *WASHC1*, *XPO4*) linked to knockout mouse models characterized by abnormal development or embryo lethality. Notably, the SNPs at *TENM4* and near *NUFIP1* were significant for the analysis of infertility. Lastly, there were two SNPs located near or at genes functionally linked to impaired fertility (*KPNA1*, *EGFR*).

**Fig. 2.**
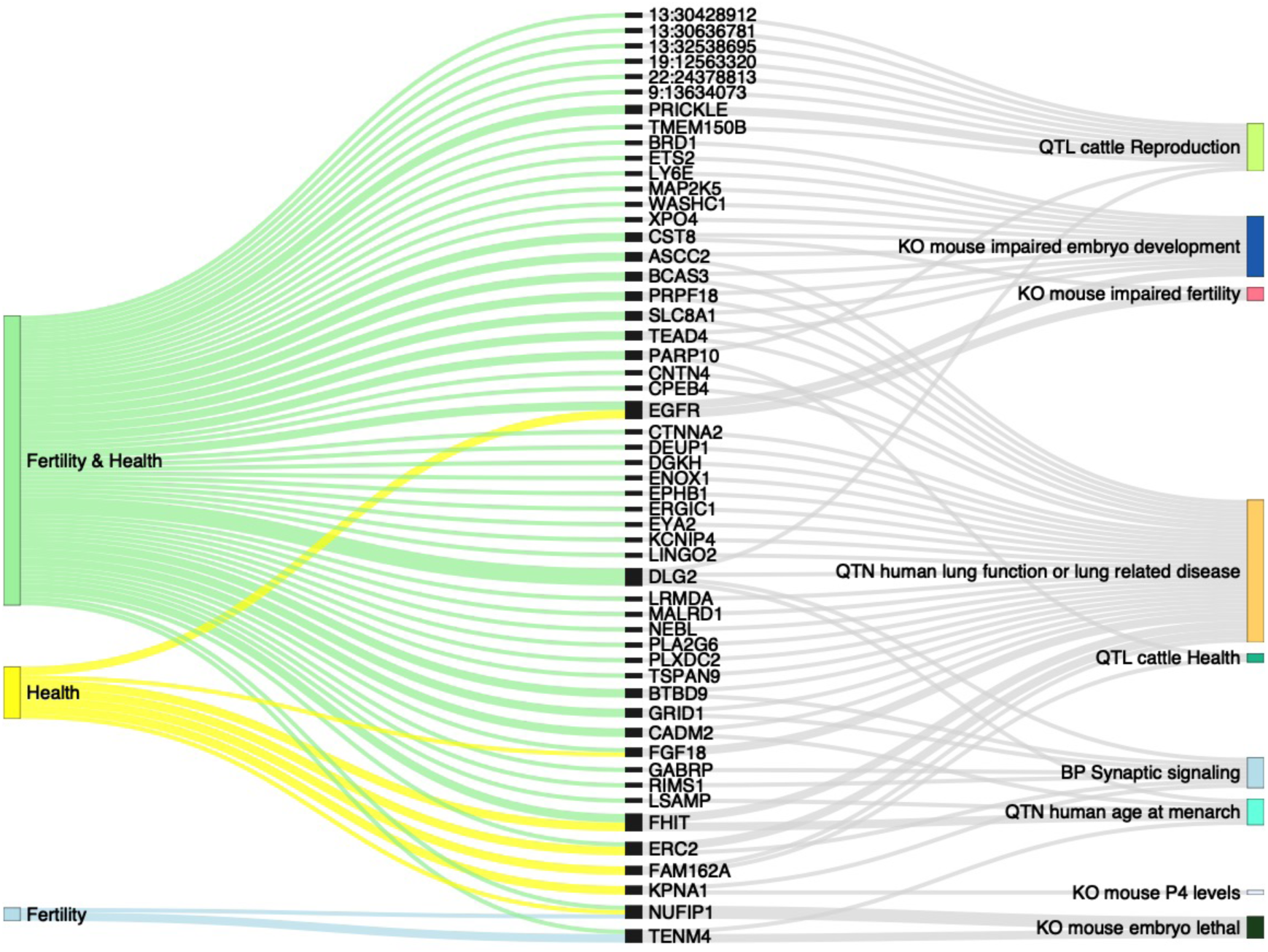
Graphical display of the *in silico* annotation of genes and SNPs that are not within or near gene boundaries. Left column represents the studies, center column indicates the genes used for annotation, and right column represents the annotations. BP: Gene Ontology Biological Process; KO: knockout, QTL: Quantitative Trait Loci, QTN: Quantitative Trait nucleotide.

We also screened annotated genes identified in our analyses using the National Human Genome Research Institute-European Bioinformatics Institute GWAS catalog [39]. In humans, polymorphisms in the genes *CADM2* [59, 60]*, DLG2* [59], *FHIT* [60], *LSAMP* [60] and *TENM4* [60] were associated with age at menarche. Also in humans, polymorphisms in the genes *ASCC2* [61], *BCAS3* [62–65], *BTBD9* [66], *CADM2* [66], *CNTN4* [66], *CPEB4* [61], *CTNNA2* [65], *DEUP1* [67], *DGKH* [66, 68], *DLG2* [66], *ENOX1* [65], *EPHB1* [69], *ERC2*[64–66], *ERGIC1* [65], *EYA2* [62, 70–72], *FAM162A* [62], *FGF18* [72], *FHIT* [73], *GRID1* [74], *KCNIP4* [66], *LINGO2* [65], *LRMDA* [62, 64, 65, 75, 76], *MALRD1* [77], *NEBL* [62, 65], *PLA2G6* [78], *PLXDC2* [62, 65], *PRPF18* [66], *SLC8A1* [65], *TEAD4* [65], *TSPAN9* [62, 65] were associated with lung function or pulmonary diseases.

### Impact of editing NUFIP1 and TENM4 genomic sequences on pre-implantation development

Considering that significant SNPs for the infertility test were near or within gene boundaries of *NUFIP1* (Fig. 3A) and *TENM4* (Fig. 3B) genes, respectively, and these genes also are associated with embryonic lethality in mouse knockouts, we interrogated whether those genes are transcribed in cattle pre-implantation embryos. Both genes are expressed in cattle oocytes and pre-implantation embryos, including blastocysts (Fig. 3C, from GSE99678, GSE199210, GSM7054290 [79–81]). While transcripts of *TENM4* are present in inner cell mass and trophectoderm in equivalent abundances ([82, 83] Fig. 3D from GSE99678 [82]), one report showed transcripts of *NUFIP1* more abundant in trophectoderm relative to inner cell mass (Fig. 3D, from GSE99678 [82]), but this difference has yet to be confirmed [83].

**Fig. 3.**
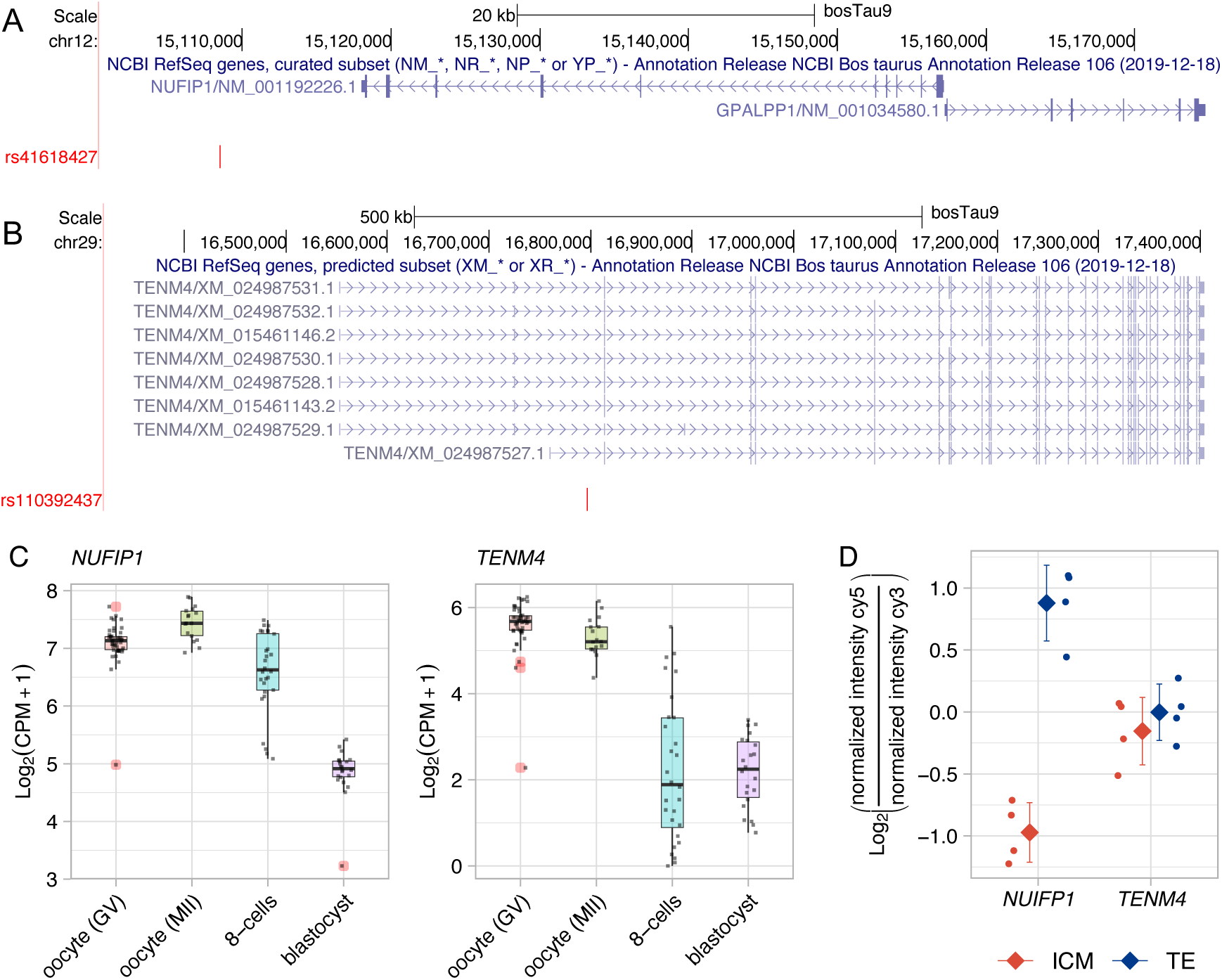
SNPs near or within boundaries of genes with knockout phenotypes leading to embryonic lethality. Position of (A) rs41618427 relative to *NUFIP1* and (B) rs110392437 relative to *TENM4*. Both images were obtained from the UCSC genome browser based on bosTau9 assembly. (C) Transcript abundances of *NUFIP1* and *TENM4* in cattle oocytes and pre-implantation embryos. Black dots are the data points, and red circles indicate outliers. Data from GSE99678, GSE199210, GSM7054290 [79–81]. (D) Relative transcript abundance of *NUFIP1* and *TENM4* in inner cell mass and trophectoderm. Dots represent the data points, diamonds represent the mean, and vertical bars represent the standard deviation of the mean. ICM: inner cell mass, TE: trophectoderm. Data from GSE99678 [82] and GSE99678 [82].

We evaluated the guide RNAs and the editing efficiency by carrying out a PCR and sequencing of the targeted DNA region. We sequenced PCR products for the region of the gene *NUFIP1* obtained from 19 embryos collected between the 8-cell stage and morula (before the effects of editing had a biological impact). Fourteen embryos tested presented both alleles edited (examples in Figure 4A), while five embryos were heterozygote presenting edited and wild-type alleles, which yields 73.7% efficiency of producing edited sequences, altering the *NUFIP1* gene on both chromosomes of treated embryos. We also sequenced PCR products for the target region of the gene *TENM4* obtained from eight embryos (three 8-cells ad five blastocysts). Seven embryos tested presented both alleles edited which yields 87.5% efficiency of producing edited sequences altering the gene on both chromosomes of treated embryos (examples in Figure 4B).

**Fig. 4.**
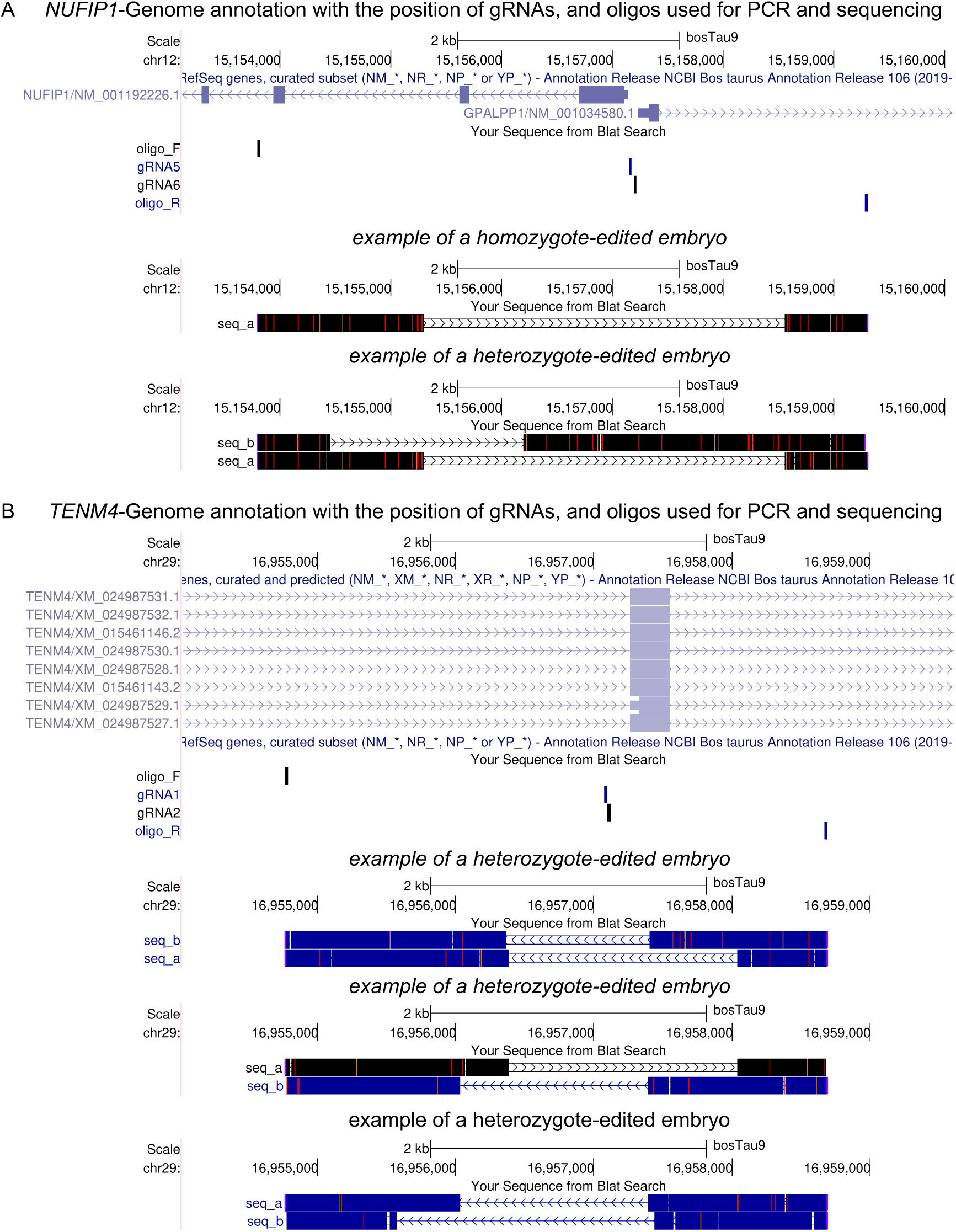
Targeted DNA editing in pre-implantation embryos. (A) Position of gRNAs and oligonucleotides used for targeted sequencing as well as examples of sequences most commonly produced in tests of the gene *NUFIP1*. (B) Position of gRNAs and oligonucleotides used for targeted sequencing as well as examples of sequences most commonly produced in tests of the gene *TENM4*.

The removal of the promotor and transcript starting site of the *NUFIP1* gene did not impact initial cleavages, evaluated at 48hpf (Fig. 5A, Supplemental Table S1-S3). By comparison, the development of blastocysts was significantly reduced when presumptive zygotes were electroporated with ribonucleoproteins targeting the *NUFIP1* gene (P<0.05, Fig. 5A, Supplemental Table S1-S3). On the other hand, deletions on the first coding exon along with the most likely translation starting site of the *TENM4* gene did not significantly reduce the ratio of putative zygotes cultured that reached blastocyst (P>0.05, 167, and 190 hpf, Fig. 5B, Supplemental Table S4-S6).

**Fig. 5.**
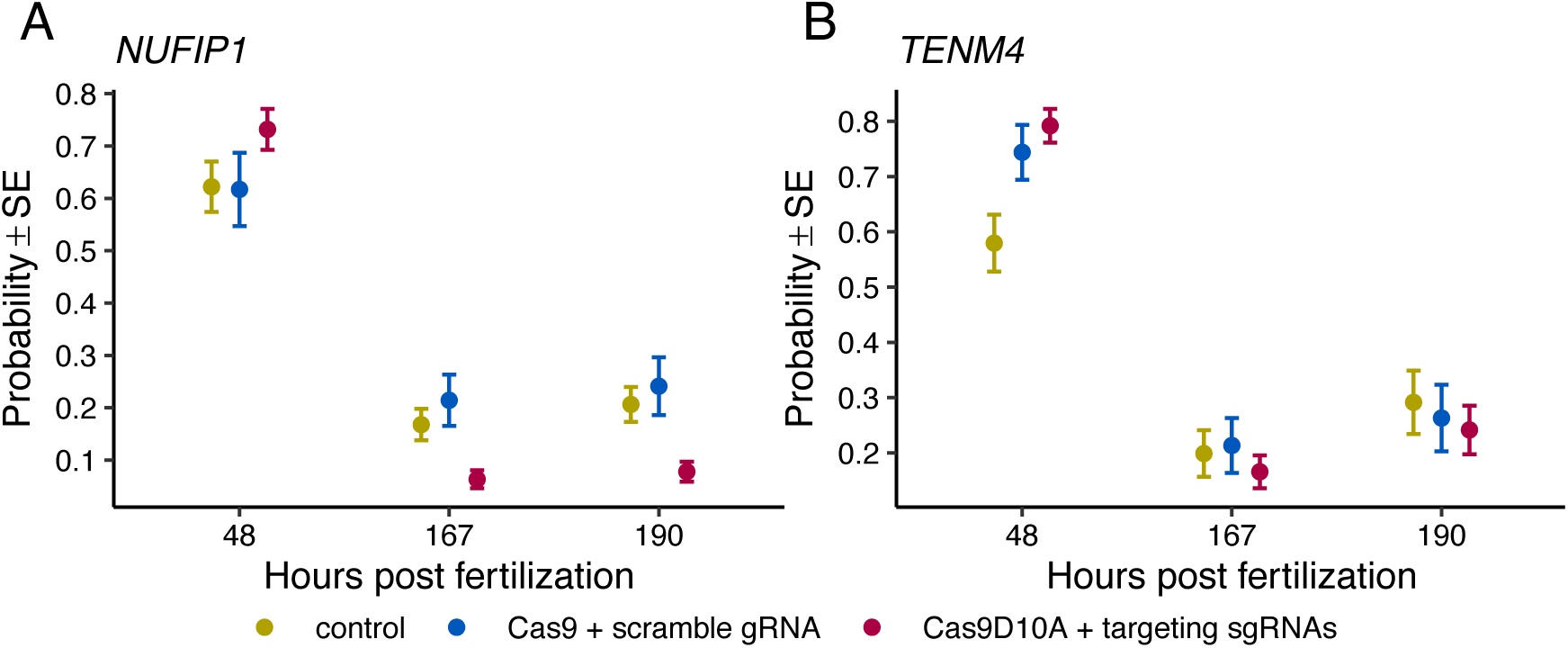
Evaluation of embryo development. The adjusted probability of embryo development at three-time points when testing the effects of deleting DNA segments of (A) *NUFIP1* and (B) *TENM4*. SE: standard error.

## Discussion

Disease and infertility are two of the most critical reasons dairy cattle leave the herd, thus posing barriers to profitable and sustainable dairy production. Genome-wide association studies have become a valuable approach to identifying genomic variants that are associated with traits related to fertility and health, and they have the potential to assist producers in making management decisions. In this study, we aimed to identify variants that were associated with health and fertility in Holstein heifers. The findings pose strong evidence that infertility in otherwise healthy heifers may be related to early embryonic losses. Furthermore, the results point to the direction that there is a genetic architecture around heifer susceptibility to diseases.

Our experimental design focused on contrasting the genetic background of groups of animals representing extreme phenotypes. While case-control designs are an effective means to identify the genetic architecture of complex traits [84], the consequence of working with individuals that represent extreme phenotypes is the limited sample size, thus limiting power [84]. The sample size is limited in the current study and certainly prevented the identification of other polymorphisms that might be of importance for the traits evaluated. Therefore, the results presented in Tables 1-3 are not an exhaustive list of polymorphisms that may be associated with health and fertility in dairy heifers. Although limited, we were able to support many of the genes with independent data from reputable databases (Animal Quantitative Trait Loci (QTL) Database [33, 34], Mouse Genome Database [37, 38], and human GWAS studies [39]) and one gene was supported by a loss of function experiment.

Our analyses identified SNPs associated with infertility and or sub-infertility, as well as with their overall health. None of the SNPs presented here have been identified in past research [2, 12–19]. One possible reason for the discrepancy is that our model compares samples from extreme phenotypes (case-control design), which contrasts with heifer conception rate, antral follicle count, and age at first calving. The other possibility is that the genetic background and limited sample size of this study represent a small subgroup of animals compared to the diverse genetic backgrounds across the US.

Our analysis revealed five SNPs associated with heifer infertility and eight SNPs associated with the number of artificial inseminations needed to deliver a calf. Notably, two SNPs were in or near genes with knockout mouse models that cause embryo lethality (*TENM4*, *NUFIP1*). The identification of rs41618427 and rs110392437 adds to a list of other SNPs [2, 12, 13, 15, 20, 85] located in or near genes previously implicated in embryo lethality in mouse knockout models. Our experiments determined that impairing the function of *NUFIP1* reduced pre-implantation embryo development. Although the ablation of *NUFIP1* does not present causality of rs110392437, it provides further evidence that some alleles have a negative impact on the reproductive performance of carrier heifers without necessarily influencing their anatomy or physiology.

A few SNPs stood out in the analysis comparing heifers that died or were culled due to diseases relative to healthy heifers. First, the SNP rs29019349 in the gene FAM162A is within QTL_IDs 159986 and 159987, which were previously associated with susceptibility to bovine respiratory disease in cattle [57]. In humans, polymorphisms in the genes *ERC2* [62, 65, 66], *FAM162A* [62] and *FGF18* [72] were associated with impaired lung function. Also, in humans, a polymorphism in the gene PAM [65] was associated with white blood cell count. Our results support previous findings that there is a genetic component to disease predisposition in cattle and aid the identification of variants that contribute to this phenotype.

The genetic architecture of female fertility is complex and intersects with other critical functions or systems, such as the immune system [17, 86]. Three observations in our results support this idea. First, several polymorphisms evidenced in the independent analyses (fertility or health) were also significant when we analyzed samples of infertile and culled heifers as one group. Second, as expected, a greater sample size adds power to the analysis, and thus, there were more statistically significant genes. Third, many genes with significant SNPs in the joint analysis (Table 3 and Figure 3) were previously associated with reproductive functions (*BRD1*, *CST8*, *EGFR*, *ETS2*, *LSAMP*, *LY6E*, *MAP2K5*, *NUFIP1*, *PRICKLE*, *TENM4*, *TMEM150B*, *WASHC1*, *XPO4*), abnormal respiratory function (*BTBD9*, *CNTN4*, *CPEB4*, *CTNNA2*, *DEUP1*, *DGKH*, *ENOX1*, *EPHB1*, *ERC2*, *ERGIC1*, *EYA2*, *FGF18*, *GRID1*, *KCNIP4*, *LINGO2*, *LRMDA*, *MALRD1*, *NEBL*, *PLA2G6*, *PLXDC2*, *TSPAN9*), or both (*ASCC2*, *BCAS3*, *CADM2*, *DLG2*, *FHIT*, *PARP10*, *PRPF18*, *SLC8A1*, TEAD4). It was also notable that the genes *BCAS3*, *EPHB1*, and *KCNIP1*, identified in our combined fertility and health study, were associated with heifer fertility traits in a recent study by Forutan et al.[87].

We investigated the genetic variation in dairy heifers that have reproductive limitations or presented irreversible health problems in comparison to dairy heifers that represent healthy and fertile heifers. The results confirm a genetic component of fertility and health. Most importantly, we show a shared genetic architecture between fertility and health. Several of the genes containing or near significant SNPs overlapped with significant associations with similar phenotypes in cattle, mouse and humans, highlighting conserved genetic elements of fertility and health across mammals.

## Acknowledgments

The authors are grateful for the collaboration with the Council on Dairy Cattle Breeding (CDCB) for extracting and releasing genotypes analyzed in this study.

## Author contributions

MM contributed to the analysis and first draft of the document and tables; GPS contributed to the processing and analysis of sequencing data; MM and AW contributed with embryo experiments; RC contributed with funding acquisition; FB obtained funding, contributed with embryo experiments, analyzed and interpreted data, analyzed and interpreted sequencing data, prepared figures and wrote the final document.

